# Density-dependent changes in neophobia and stress-coping styles in the world’s oldest farmed fish

**DOI:** 10.1101/394288

**Authors:** T. Champneys, G. Castaldo, S. Consuegra, Garcia de Leaniz

## Abstract

Farmed fish are typically reared at densities much higher than those observed in the wild, but to what extent crowding results in abnormal behaviours that can impact welfare and stress coping styles is subject to debate. Neophobia (i.e. fear of the ‘new’) is thought to be adaptive under natural conditions by limiting risks, but it is potentially maladapted in captivity, where there are no predators or novel foods. We reared juvenile Nile tilapia (*Oreochromis niloticus*) for six weeks at either high (50g/L) or low density (14g/L), assessed the extent of skin and eye darkening (two proxies of chronic stress), and exposed them to a novel object in an open-test arena, with and without cover, to assess the effects of density on neophobia and stress coping styles. Fish reared at high density were darker, more neophobic, less aggressive, less mobile and less likely to take risks than those reared at low density, and these effects were exacerbated when no cover was available. Thus, the reactive coping style shown by fish at high density was very different from the proactive coping style shown by fish at low density. Our findings provide novel insights into the plasticity of fish behaviour and the effects of aquaculture intensification on one of the world’s oldest farmed and most invasive fish, and highlight the importance of considering context. Crowding could have a positive effect on the welfare of tilapia by reducing aggressive behaviour, but it can also make fish chronically stressed and more fearful, which could make them less invasive.

## Introduction

Food security relies on Aquaculture intensification to maximise fish production and increase economic viability (d’Orbcastel et al., 2010; Suresh & Lin, 1992) but to what extent crowding may affect behaviours and compromise welfare is unclear. Stocking density is one of the parameters most easily managed by fish farmers, but what constitutes an acceptable stocking density differs from species to species, and is subject to debate (Conte, 2004; Ellis et al., 2002; Felicity A Huntingford et al., 2006; Turnbull et al., 2008). For example, experiments on Atlantic salmon (*Salmo salar* L) and African catfish (*Clarias gariepinus*) indicate that fish reared at high density become less aggressive but also more willing to rise to the surface to feed than fish reared at low density (Fenderson & Carpenter, 1971; van de Nieuwegiessen, Ramli, Knegtel, Verreth, & Schrama, 2010), but the opposite has also been reported, with fish becoming more aggressive as stocking density increases (Rueda, 2004). Even within siblings, stress-related behaviours may increase with increasing density in some individuals but decrease in others (Conte, 2004).

(FAWC, 2005) recommends that fish are given ‘sufficient space to show most normal behaviours with minimal pain, stress and fear’, but few studies have addressed what constitutes ‘normal behaviours’, what is ‘sufficient’ space, or how to detect ‘fear’ under aquaculture conditions. There is, nevertheless, consensus that an incomplete repertoire of natural behaviours may be indicative of compromised welfare (Melfi & Feistner, 2002). Unlike other behaviours, such as aggression and boldness, which have been well studied in many species (e.g. Nile tilapia, (Mesquita, Torres, & Luz, 2016); salmonids: (L.J. Roberts, Taylor, Gough, Forman, & Garcia de Leaniz, 2014); (Stringwell et al., 2014) zebrafish, (Wright, Rimmer, Pritchard, Butlin, & Krause, 2003); poeciliids, (C. Brown, Jones, & Braithwaite, 2005)), studies of fear and neophobia in fish are scant (Ashley & Sneddon, 2008; Elvidge, Chuard, & Brown, 2016). A search of 3,626 articles on fish behaviour published between 2000 and 2017 in the Journal of Fish Biology revealed only one article with the terms ‘fear’ or ‘neophobia’ (fear of the new) in the title, and only 2.5% of articles refer to fear in the text. This serves to emphasise the need for more research on fish fear, particularly on commercially important fish like Nile tilapia (*Oreochromis niloticus*), which are typically reared at very high densities, up to 100g/L (Rakocy, 2005).

When confronted with an unfamiliar stimulus many species exhibit caution and display an avoidance response (Aitken, 1972)). Such fear of the ‘new’ has been observed across taxa (Greenberg, 1990) and is most widely documented in the context of novel food and predator avoidance. According to ‘the dangerous niche hypothesis’, wariness of novel stimuli is likely to be adaptive in areas of high predation risk or when prey can be toxic (Greenberg, 2003)). By avoiding unfamiliar foods, animals can avoid harmful ones, which will often be evolutionary advantageous (Bókony, Kulcsár, Tóth, & Liker, 2012)). Neophobia can also represent an effective anti-predatory strategy (G. E. Brown, Ferrari, Elvidge, Ramnarine, & Chivers, 2013) because, unlike costly specialised defence structures (such as spines or armoured plates) that can lose their effectiveness over time (Marchinko, 2009); (Chivers, Zhao, Brown, Marchant, & Ferrari, 2008)) or become obsolete in novel environments (Leaver & Reimchen, 2012), neophobia will generally reduce the chances of suffering a predatory attack (Meuthen, Baldauf, Bakker, & Thunken, 2016). However, neophobia also has costs. For example, a persistent fear of novel stimuli can be accompanied by an acute stress response and elevated levels of corticosteroids (Coppinger, 1970); (Greenberg, 1990)), or may result in missed feeding opportunities and unnecessary energy expenditure. Neophobia, therefore, is thought to be most beneficial in high predation environments (Ferrari, McCormick, Meekan, & Chivers, 2015), but how it may evolve under predator-free aquaculture conditions is unclear. Some studies suggest that highly plastic species (i.e typically generalists) are less neophobic than specialists that have a narrower niche breadth (Webster & Lefebvre, 2000), but neophobia is also well documented among trophic opportunists that have a wide diet (Barnett, 1958); (Heinrich, 1988);(Moldlinska, Stryjek, & Pisula, 2015). The fact that neophobia is present in species that occupy the full foraging continuum serves to highlight the difficulties of explaining why variation exists, at both inter and intra-specific level (Greenberg, 1990).

Neophobia may be expected to vary with genetic background and early experience (Fox & Millam, 2004)), but how it relates to rearing density and stress coping styles is not known. Historically, tilapias have had a reputation for being tolerant of very high stocking densities (El-Sayed, 2006; Rakocy, 2005), but as the welfare of farmed fish receives more attention (Ashley, 2007; F.A. Huntingford & Kadri, 2009)) - and as behavioural metrics of welfare become more widely used (Martins et al., 2012)), this tenet is coming under closer scrutiny. Tilapias rank among the oldest, and most widely farmed fish worldwide (El-Sayed, 2006; Rakocy, 2005), but are also included in the ‘100 World’s Worst Invasive Alien Species’ (Lowe, Browne, Boudjelas, & De Poorter, 2004), so knowledge about how these species respond to novelty when they escape from fish farms and become feral might be important for reducing impacts. To address these questions, we reared juvenile Nile tilapia at high and low densities, measured the extent of eye and skin darkening (two metrics of chronic stress, (Volpato, Luchiari, Duarte, Barreto, & Ramanzini, 2003); (Vera Cruz & Tauli, 2015), and screened them for neophobia, as well as for activity, aggression, and boldness, as these are measures of strese coping style (Barreto, Carvalho, & Volpato, 2011); (Rey et al., 2016)). We used fry because this allowed us to address the effect of rearing density at a critical stage of development, when the young leave the care of their parents and variation in exploratory behaviour is first manifested (Kawamura & Washiyama, 1989). Chronic stress during early life is known to cause long-term changes in boldness and neophobia in rats (Chaby et al., 2013 (Chaby, Cavigelli, White, Wang, & Braithwaite, 2013), so our hypothesis was that changes in rearing density might also trigger similar changes in neophobia and coping style in fish.

## Materials and Methods

### Experimental design

Ten-days old, mixed-sex Nile tilapia were sourced from Fishgen and reared for six weeks at either high density (50g/L; 80 fish/tank) or low density (14g/L; 20 fish/tank) in identical 28L white opaque plastic tanks (40L x 31W x 23H cm) at CSAR’s tilapia recirculation facility. These densities fall towards the upper and lower end of densities commonly found at recirculation aquaculture systems for this species (Conte, 2004). Low density fish were reared in triplicate and high density fish in duplicate tanks. Photoperiod was maintained at 12D:12L and temperature was set at 25.2C (SE± 0.25). Fish were fed *ad libitum* (1.5mm Nutra pellets) twice a day (08:30 and 16:00 hrs). Mean weight after six weeks was 12.5g for high density fish (SE±3.7) and 14.4g (SE±4.8) for low density (t = −2.390, df = 106.96, *P* = 0.019). There were no mortalities in any of the tanks during the course of the study.

### Behavioural screening

The test arenas consisted of two plastic tanks, identical to the rearing tanks (40L x 31W x 23H cm) filled with 15L of water that were divided into four equally spaced sections: an acclimatisation section fitted with a removable cover and a sliding door that could be lifted remotely (zone 0), and three open test sections without cover (zones 1, 2 and 3) that were delineated by marks on the tank bottom. To test for neophobia, a highly conspicuous plastic green toy was fixed with silicon to section 2 (in the middle of the test arena) to serve as a novel object, as per (Brydges, Boulcott, Ellis, & Braithwaite, 2009); **Figure 1**).

**Figure 1.**
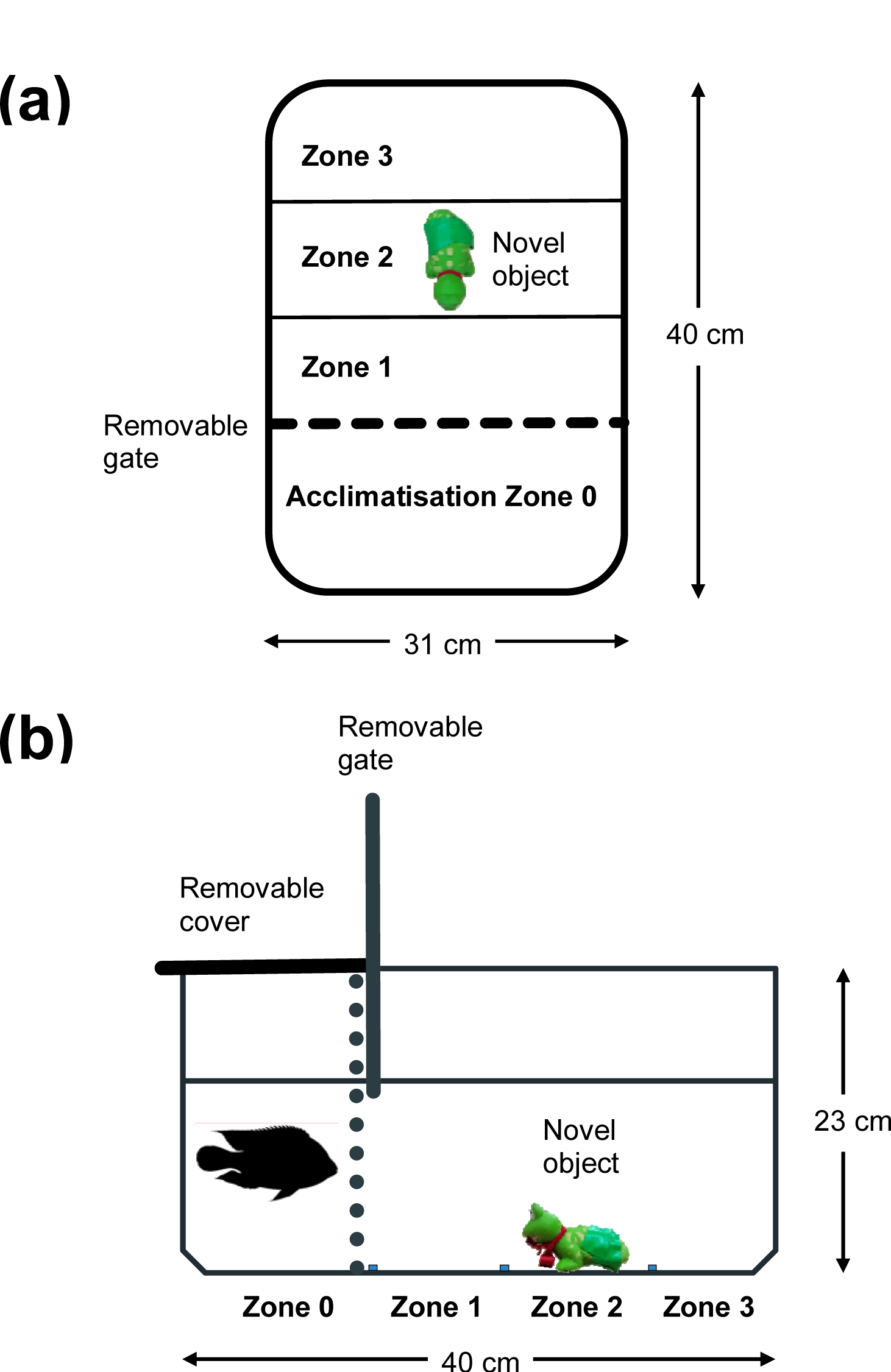
Diagram of the test arena used to assess neophobia and coping style in juvenile Nile tilapia (a, top view; b, side view – not to scale). The acclimatisation zone (Zone 0) is separated by an opaque sliding door that can be raised remotely to within 1 cm below the water surface. Zones 1, 2 and 3 are delineated by lines drawn below the tank. The novel object, a conspicuous green plastic toy, is glued to the centre of Zone 2. The overhead cover was left in place or removed, depending on test.

After six weeks of rearing, fish were netted haphazardly from each tank and placed singly into the acclimatisation chamber for 10 minutes, after which time the sliding door was slowly raised, and the fish behaviour recorded for 15 minutes with an overhead camera (GoPro Hero 5) fixed 1m above the tanks. At the end of the test period, the fish were removed, measured (1 mm), weighed (0.1 g) and the test tanks drained, rinsed, and refilled with water to remove any olfactory cues that might affect subsequent behaviours. Two blocks of trials were allocated at random and run concurrently with or without cover in the acclimatisation chamber during testing. In total 116 fish were tested in a 2×2 balanced, fully factorial design representing two rearing densities (high vs low density; n = 29 each), and two test conditions (cover vs no cover; n = 29 each). All tests took place over a five-day period (1-5 Aug 2016).

Videos were reviewed using VLC Media Player and analysed by the first author to ensure consistency. Four behaviours were quantified on each fish to assess neophobia and coping style: (1) the latency (secs) to exit the acclimatisation zone once the door was lifted (*boldness*), (2) the number of times the fish came within 2 cm of the novel object but did not touch it (*approaches*), (3) the number of times the fish charged towards the novel object (*attacks*), and (4) the time spent (secs) in each of the tank zones, from which we computed the mean distance to the novel object (*distance*), and the Shannon evenness index to measure spatial use (*activity*). We assumed that the greater the distance to, or time spent away from, the novel object, the more fearful the fish were (Brydges et al., 2009)).

### Assessment of skin and eye darkening

To assess the effect of rearing density on eye and skin darkening, tilapia fry of the same age (10-12 days) and origin as above were reared in a concurrent experiment for 10 weeks, using the same densities (high density: 80 fish/tank; low density: 20 fish/tank) and under the same conditions as per the behavioural experiment above. A sample of 107 fish (n= 52 low density; n = 55 high density) was then photographed (Canon EOS 400D Digital) inside a glass aquarium against a standard background (Classic Target–Xrite–Color checker). Colour measurement and standardization were performed using pixel sampling with GIMP 2.8.16 as per (Clarke & Schluter, 2011)). Briefly, grayscale values (0–255) of R, G and B were measured along the fish flank, in a standardised window defined by the beginning and end of the dorsal fin. As we were interested in the extent of darkening, rather than in colouration, RGB values were converted to HSV space using the *rgbtohsv* function in the *grDevices* R package and the brightness (luminance) value was used for statistical analysis (Hanbury, 2002). To quantify eye darkening, we used a modification of the method proposed by (Freitas, Negrao, Felicio, & Volpato, 2014): we divided photographs of the eye sclera into eight equal sections and counted the number of darkened sections.

### Statistical analysis

We used R version 3.4.3 for all analyses (Core Team, 2014). As stocking density was found to affect body size (fish reared at high density were smaller at the end of the experiment *t*_109.21_= −3.07, *P* = 0.003), and this could affect behaviour, body size was included as a covariate in statistical analyses.

Latency to leave shelter was tested by right censored Cox proportional hazards regression, as this can accommodate both continuous (fish length) and categorical predictors (cover and density) and can assess their joint effects simultaneously (Crawley, 2007). We used the *coxph* function in the *survival* R package (Therneau, 2018) to compute the Cox proportional hazards regression and the *survminer* package (Kassambara, Kosinski, Biecek, & Fabian, 2018) to visualize the results. Starting with a maximal model with rearing density, cover, fish length and all their interactions, we employed the *step* and *dredge* functions for variable selection to arrive at a minimal adequate model (Crawley, 2007).

The number of attacks and approaches was analysed using a Generalized Linear Mixed Effects Model (GLMM) with a Poisson link-function using the *lme4* package (Bates et al 2015 (Bates, Mächler, Bolker, & Walker, 2015)) with rearing density, cover and fish length as fixed effects, and tank identity as a random effect. The average distance to the novel object was computed from the time the fish spent in each zone, and this was analysed as a Linear Mixed Model (LMM) using *lme4* as above. Skin darkening was analysed as a function of ‘luminance’ (brightness) via LMM using density, body weight and sex as fixed effects and tank identity as a random effect. Model simplification was achieved by starting with a maximal model with all main effects and interactions and arriving at a minimal adequate model with comparisons made by Maximum Likelihood on the basis of AIC values and the *anova* command; the final model was refitted by Restricted Maximum Likelihood (Crawley, 2007).

### Ethics

The work carried out was screened and approved by the College of Science Ethics Committee on 09/06/2016.

## Results

### Latency to leave shelter

Latency to leave shelter depended on rearing density (*z* = 2.22, *P* =0.027), the presence of cover during testing (*z* = −5.13, *P*<0.001) and the interaction between density and cover (z = 2.15, *P* = 0.031). Neither body size, nor any other interactions, were significant (overall model fit LRT =75.2, df = 3, *P* < 0.001 for n= 116 and n= 77 events). Fish reared at high density were more cautious and less likely to leave the starting box (only 17% left) than fish at low density (50% left), but as expected, this was only manifested when the starting box remained covered during testing (Cox estimate for low density = 2.06, SE = 0.55, *z* = 3.73, *P* <0.001; n= 58, number of events= 24, LRT = 19.59, df = 1, *P*<0.001; **Figure 2a**). When the starting box was left uncovered (and thus no shelter was afforded), nearly all fish exited the starting box (53/58 or 91%), regardless of rearing density (**Figure 2b;** Cox estimate for low density = 0.52, SE = 0.28, *z* = 1.87, *P* =0.061; n= 58, number of events= 53, LRT = 3.51, df = 1, *P*=0.061).

**Figure 2.**
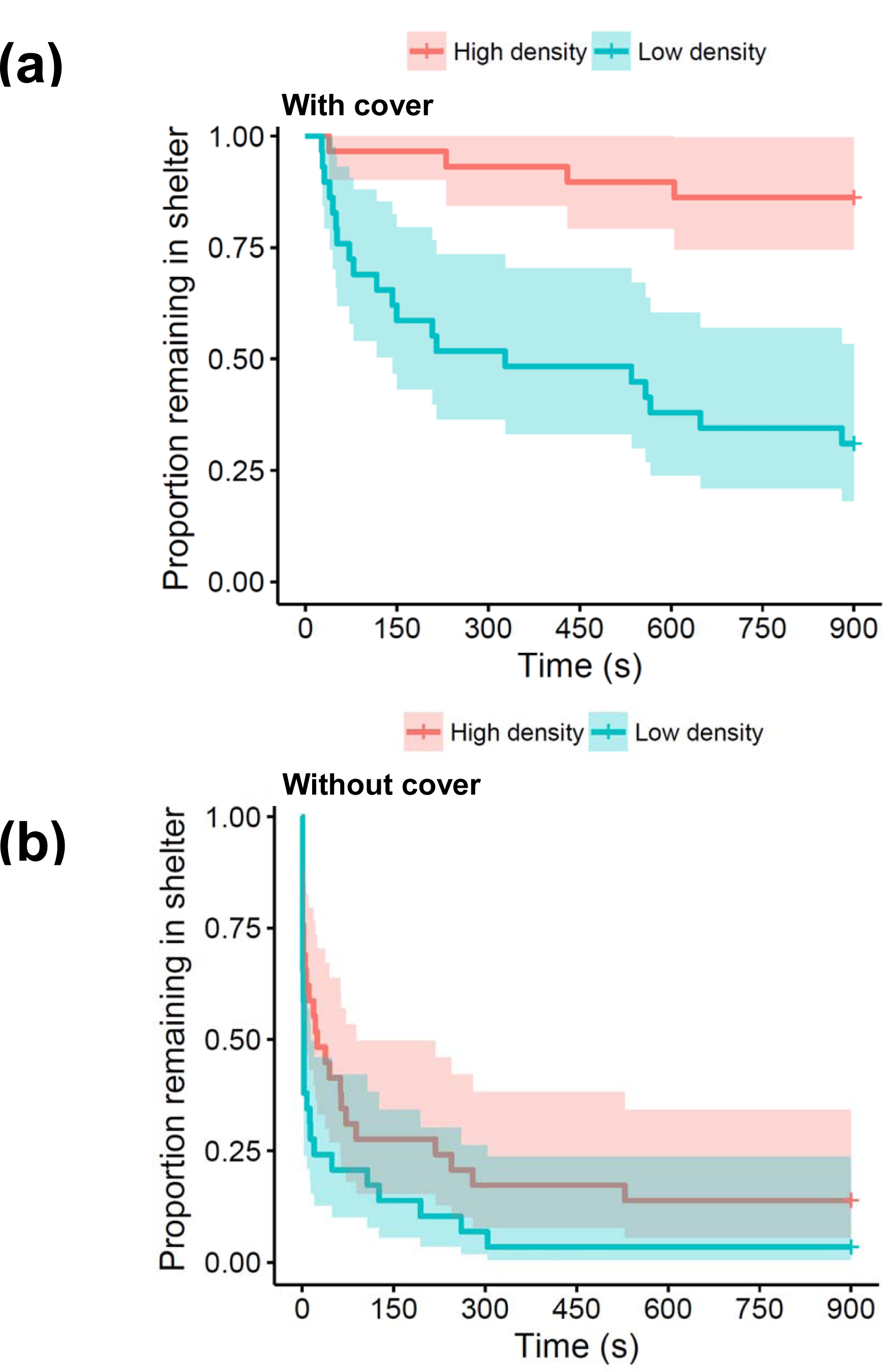
Latency to leave shelter (s, ± 95 CI) of juvenile Nile tilapia reared at low (14 g/L) and high (50 g/L) density with (a) and without (b) overhead cover in the acclimatisation area.

### Number of approaches

The number of approaches to the novel object depended mostly on rearing density (fish at low density made significantly more approaches than fish at high density (GLMM estimate = 2.04, SE = 0.26, z-value = 7.854, *P* < 0.001), and to a lesser extent on the presence of cover (fish provided with cover made fewer approaches, estimate = −1.19, SE = 0.15, z-value = − 7.48, *P* < 0.001), body size (larger fish made more approaches (estimate = 0.19, SE = 0.07, z-value = 2.66, *P* = 0.008), and the interaction between cover and body size (estimate = −0.39, SE = 0.14, z-value = −2.81, *P* = 0.005).

### Number of attacks

As with the number of approaches, the number of attacks was significantly higher among fish reared at high density than at low density (estimate = 2.20, SE= 0.43, z-value = 5.11, *P* < 0.001), and also higher among fish tested without cover than with cover (estimate = −1.87, SE = 0.38, z-value = −4.93, *P* < 0.001). None of the interactions were significant.

### Neophobia

The distance that fish mantained to the novel object depended on the density they had been reared at, as well as the presence of cover (**Figure 3**). Fish reared at high density were more neophobic, i.e. stayed further away from the novel object, than fish reared at low density (estimate for high density = 1.17, SE = 0.37, *t*_113_ = 3.14, *P* = 0.002). Likewise, fish tested with cover were also more neophobic than fish tested without cover (estimate = 4.39, SE = 0.37, *t*_113_ = 11.8, *P* < 0.001). None of the interactions were significant. Overall, fish reared at low density spent more than twice as long in the vicinity of the novel object (i.e. Zone 3) than fish at low density (mean time ± SE low density = 94.0±13.5 s; mean time ± SE high density = 44.34±13.5 s; *t*_114_ = 2.60, *P* = 0.010).

**Figure 3.**
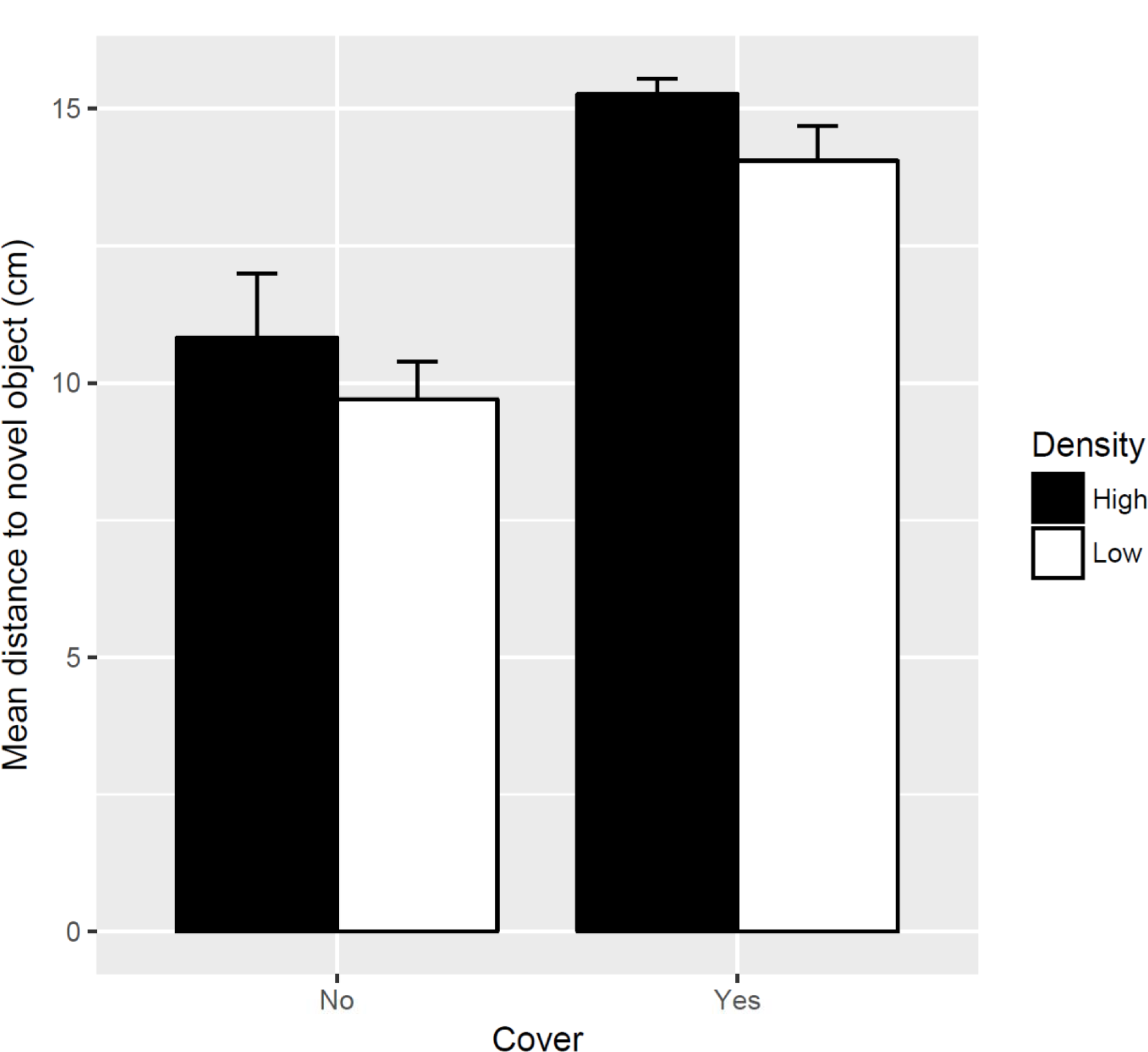
Distance to a novel object (mean ± 2 SE) maintained by juvenile Nile tilapia reared at low (14 g/L) and high (50 g/L) density.

### Skin and eye darkening

Fish reared at high density were significantly darker (i.e had lower values of luminance) than fish reared at low density (**Figure 4;** estimate = −0.072, SE = 0.017, *t*_105_ = −4.29, *P* < 0.001), while body size and sex had no effect. Eye darkening was positively correlated with skin darkening (*r* = 0.28, *t*_105_ = 2.98, *P* = 0.003), suggesting that both metrics responded in the same way to rearing density.

**Figure 4.**
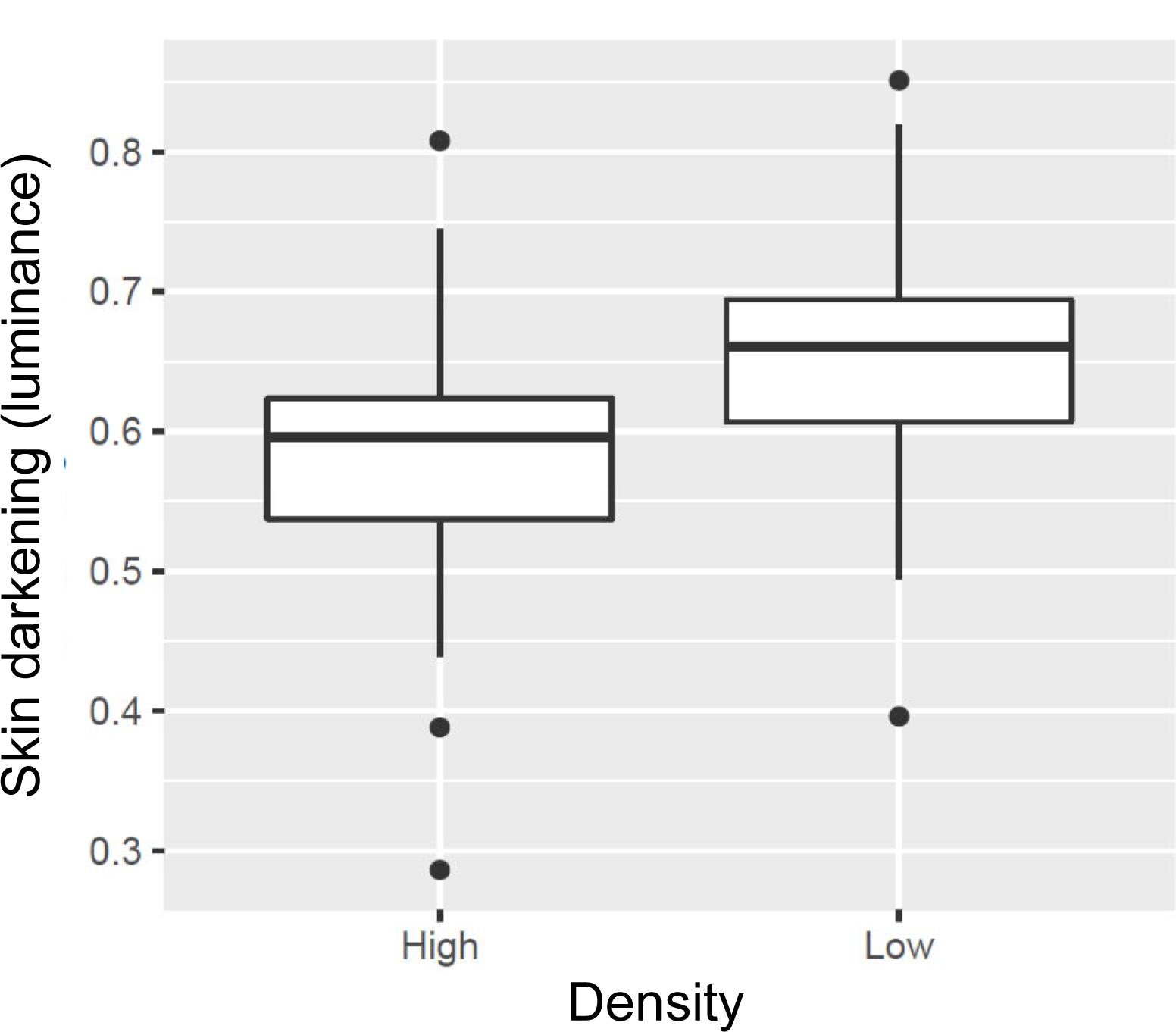
Skin darkening (luminance) of juvenile Nile tilapia reared at low (14 g/L) and high (50 g/L) density.

## Discussion

Freedom from fear and freedom to express normal behaviours are two of the five metrics of good animal welfare (FAWC, 2005), but no studies have assessed how rearing conditions affect fear in farmed fish. Our study indicates that rearing density has a strong effect on neophobia (fear of the new) and stress-coping styles in Nile tilapia, one of the world’s oldest and most widely farmed fish (El-Sayed, 2006). Fish reared at high density were darker, more neophobic, less aggressive, less mobile, and less likely to take risks than those reared at low density, and these effects were exacerbated when no cover was available. Thus, tilapia reared at high density displayed a reactive stress coping style (*sensu* (Koolhaas et al., 1999); (Coppens, de Boer, & Koolhaas, 2010) while those reared at low density displayed a proactive style (**Table 1**).

**Table 1.** Effects of rearing density on neophobia and coping style in juvenile Nile tilapia screened singly without cover (means ± SE).

| Trait | Metric | Rearing Density |  |
| --- | --- | --- | --- |
|  |  | Low<br>(14g/L) | High<br>(50 g/L) |
| Neophobia | Distance to novel object (cm) | 9.7±0.35<br>close | 10.8±0.58<br>far |
| Eye darkening | Sclera score (0-8) | 1.15±0.21<br>pale | 1.29±0.17<br>dark |
| Skin darkening | Luminance | 0.65±0.01<br>pale | 0.58±0.01<br>dark |
| Activity | Evenness of tank use (0-2) | 1.47±0.11<br>active | 0.84±0.12<br>sedentary |
| Aggression | No. attacks | 1.62±0.39<br>aggressive | 0.17±0.07<br>fearful |
| Exploratory behaviour | No. approaches | 5.90±0.81<br>exploratory | 0.79±0.24<br>cautious |
| Risk-taking | Latency to leave shelter (s) | 69.9±33.8<br>bold | 184.3±59.2<br>shy |
| Coping style |  | *Proactive* | *Reactive* |

A reactive coping style tends to be associated with shyness, sedentarism, low sympathetic reactivity and high hypothalamus-pituitary-adrenal (HPA) axis activity (Koolhaas et al., 1999; Korte, Koolhaas, Wingfield, & McEwen, 2005) and a recent review suggests that these traits may be common among fish that experience sustained chronic stress in aquaculture (Castanheira et al., 2017). Our study suggests that neophobia is part of a reactive coping style, and that this also includes body and eye darkening. Neophobia is expected to increase in dangerous environments (Greenberg, 2003), and should therefore be maladaptive in captivity because there is no risk of predation, and the probability of encountering dangerous foods is low. Yet, a meta-analysis shows that captive bred individuals tend to be more neophobic than wild ones (Crane & Ferrari, 2017), suggesting that there are other considerations at play. An incomplete repertoire of natural behaviours may be indicative of compromised welfare in captive animals (Melfi & Feistner, 2002), but behaviours that are adaptive in the wild may become maladaptive in captivity, and vice-versa (Stringwell et al., 2014). Likewise, while fear and stress may play an adaptive role in natural populations, they are generally considered undesirable in an aquaculture setting (F. A. Huntingford, 2004; Felicity A Huntingford et al., 2006) (Ashley, 2007)).

Hatchery-reared fish tend to show increased boldness and aggression (typical of a proactive coping style) compared to wild fish (Stringwell et al., 2014), (L.J. Roberts, Taylor, & Garcia de Leaniz, 2011); (L.J. Roberts et al., 2014), but this may not necessarily result from high densities per se, but rather from selective breeding for fast growth, scramble for food, and absence of predators in artificial environments ((Felicity A Huntingford et al., 2006)). Neophobia has been associated with stress and a reactive coping style in roe deer (Monestier et al., 2017), and also with stress and poor welfare in aquaculture (Martins et al., 2012). For example, Mozambique tilapia reared in socially unstable groups (and thus subject to social stress) show increased neophobia (Galhardo, 2010), and high density in our study also resulted in more neophobic fish. This suggests that crowding, neophobia, and stress are likely related in Nile tilapia.

We did not measure stress-related hormones in our study, but high densities have previously been found to increase plasma cortisol in Nile tilapia, resulting in weight loss and a heightened response to subsequent acute stressors (Barcellos, Nicolaiewsky, De Souza, & Lulhier, 1999); (El-Sayed, 2002). Thus, the positive association we found between high density and body and eye darkening (as well as a reduced weight compared to low density fish) suggests that darkening is probably a stress response in Nile tilapia. Skin darkening in teleosts is mediated by changes in the motility and density of melanophores, both of which can change rapidly under hormonal control, and in response to rearing conditions ((Michalis Pavlidis, Karkana, Fanouraki, & Papandroulakis, 2008). Melanocyte hormone stimulates (α-MSH) and melanin concentrates hormone (MCH) are the two main hormones involved in the modulation of chromatophore dispersal (Höglund, Balm, & Winberg, 2000); (Fujii, 1993, 2000) (Burton, 2002)), and it has been shown that social stress can increase α-MSH in plasma and result in body darkening in salmonids (Gilham & Baker, 1985; Green & Baker, 1991). Eye and body darkening are associated with subordinance (Ramanzini, Volpato, & Visconti, 2018; Volpato et al., 2003) and a reactive coping style in Nile tilapia (Vera Cruz & Tauli, 2015), as found in our study. Thus, our results are consistent with the idea that crowding during aquaculture intensification makes Nile tilapia chronically stressed, and as seen in other species, this results in body and eye darkening, neophobia, and - more generally, in a shift from a proactive to a reactive stress coping style.

Central to the concept of allostasis is that stress coping styles are consistent over time and across situations (Coppens et al., 2010); (Gosling & John, 1999), being synonymous with behavioural syndromes, temperaments and personalities (Koolhaas et al., 1999); (Coleman & Wilson, 1998)). These tend to map well into the bold-shy continuum, the dominant vs. subordinate behaviours, the aggressive vs. submissive response, and the hawk vs. dove strategies (Coppens et al., 2010; Koolhaas et al., 1999). However, in our study stress-coping styles varied depending on rearing conditions, and were also context-dependent, as shown by the contrasting effects of overhead cover. When overhead cover was in place during testing, the effects of rearing density on coping styles were greatly diminished. It was only when cover was removed, and no hiding place was provided, that a strong difference in coping styles became apparent between rearing densities. A similar situation has been reported with regards to hypoxia, with stress coping styles only becoming manifested when fish were tested under low oxygen conditions (Laursen, Olsén, Ruiz-Gomez, Winberg, & Höglund, 2011); (Killen, Marras, Ryan, Domenici, & McKenzie, 2012)). This serves to highlight the fact that fish are sufficiently plastic to adjust their behaviour in a context-dependent way to achieve allostasis, i.e. to maintain stability through change (Korte et al., 2005).

## Conclusions and prospects

To make the activity economically viable, intensive fish farming relies in growing fish at unnaturally high densities (Costa-Pierce, 2002) and our study provides novel insights into the effects of aquaculture intensification on neophobia and stress coping styles in one of the most widely farmed fish. In birds, neophobia is shaped at the chick stage (Fox & Millam, 2004), and in our study the behavioural effects of rearing density took place soon after fry left the safety of the mother’s buccal cavity, coinciding with the differentiation of the sensory system on this species (Kawamura & Washiyama, 1989). Early rearing conditions have a marked effect on brain biochemistry, catecholaminergic signalling and patterns of gene expression in zebrafish (Michail Pavlidis, Theodoridi, & Tsalafouta, 2015) and gilthead sea-bream (Vindas et al., 2018), and the same probably happens in Nile tilapia. As fry in our study had a uniform genetic background and there was no mortality during the experiment (so we can rule out selection), it seems likely that the effects of rearing density were mediated through changes in gene expression, and this warrants further study.

The results of our study could have implications for welfare and management. For example, skin and eye darkening appear to be related to stress coping styles, and given the relative simplicity of measurement, these could be incorporated into operational metrics of fish welfare, applicable under aquaculture conditions. Our results could also have implications for invasion biology because conditions that promote neophobia and a reactive coping style are expected to decrease invasion success (Conrad, Weinersmith, Brodin, Saltz, & Sih, 2011). Thus, it might be possible to reduce the invasiveness of species like Nile tilapia, which have been translocated all over the world and pose a major threat to native biodiversity (Canonico, Arthington, McCrary, & Thieme, 2005). In many species, invasive individuals often display increased activity, aggression, and boldness, traits that have been termed an “invasion syndrome” ((Merrick & Koprowski, 2017), and that our study shows are affected by rearing density, at least in Nile tilapia. Tilapia that display a reactive coping style take longer to navigate through a maze (Mesquita et al., 2016), as do guppies reared at high density (Chapman, Ward, & Krause, 2008). This suggests that reactive fish may have impaired cognition and learning, which along with reduced activity and shyness, could make them less successful at invading novel habitats. Given the threat that non-native fish pose for global biodiversity, and the major role that aquaculture plays in the introduction of invasive fish (e.g. (Garcia de Leaniz, Gajardo, & Consuegra, 2010) (Consuegra, Phillips, Gajardo, & Garcia de Leaniz, 2011), the potential for manipulating or selectively breeding fish with reactive coping styles to decrease invasiveness merits further attention.

## Acknowledgments

This work was partially funded by NRN-LCEE AquaWales Project. We are grateful to Dr.R. Stringwell, P.N. Howes and the staff at CSAR for help setting up the experiments, and to Dr. D. Gonzalez-Barreto for help with body colour measurements.

## Authors’ contributions

SC and CGL wrote the grant and secured the funding. TC, CGL and SC designed the study. TC and GC collected the data. TC and CGL wrote the MS with input from GC and SC. TC and CGL carried out the statistical analysis. All authors approved the final submission.

